# Breaking the Deadlock of Molecular Chaperones

**DOI:** 10.1101/176693

**Authors:** Tania Morán Luengo, Roman Kityk, Matthias P. Mayer, Stefan G. D. Rüdiger

**Affiliations:** Cellular Protein Chemistry, Bijvoet Center for Biomolecular Research, Utrecht University, Padualaan 8, 3584 CH Utrecht, The Netherlands; Science for Life, Utrecht University, Padualaan 8, 3584 CH Utrecht, The Netherlands; Zentrum der Molekularen Biologie der Universität Heidelberg (ZMBH), DKFZ-ZMBH-Alliance, Im Neuenheimer Feld 282, 69120 Heidelberg, Germany

## Abstract

Protein folding in the cell requires ATP-driven chaperone machines. It is poorly understood, however, how these machines fold proteins. Here we propose that the conserved Hsp70 and Hsp90 chaperones support formation of the folding nucleus by providing a gradient of decreasing hydrophobicity. Early on the folding pathway Hsp70 uses its highly hydrophobic binding pocket to recover a stalled, unproductive folding intermediate. The aggressive nature of Hsp70 action, however, blocks productive folding by grabbing hydrophobic, core-forming segments. This precludes on-pathway nucleation at high, physiological Hsp70 levels. Transfer to the less hydrophobic Hsp90 enables the intermediate to resume forming its folding nucleus. Subsequently, the protein enters a spontaneous folding trajectory towards its native state, independent of the ATPase activities of both Hsp70 and Hsp90. Our findings provide a general mechanistic concept for chaperoned protein folding.

## Introduction

It is the primary sequence of the protein that determines its native fold ^1^. Proteins condense around an initial nucleus to ultimately shape up to always the same structure ^2^. In the cell, conserved families of molecular chaperones support folding of their proteins in an energy-consuming manner, presumably by repeated cycles of binding and release ^3–6^. The molecular determinants of assisted protein folding, however, remain largely enigmatic. Here we describe that the ubiquitous, ATP-dependent chaperone machines Hsp70 and Hsp90 form a conserved cascade that promotes spontaneous protein folding by a stop-start mechanism.

Hsp90 acts downstream of Hsp70, but its contribution to protein folding is unclear ^7–10^. Under certain conditions Hsp70 can refold proteins in the absence of Hsp90 ^11^. However, molecular understanding of the Hsp70 folding mechanism requires solving two paradoxes: (i) Hsp70 binds to short, very hydrophobic stretches, which are required to build the hydrophobic core ^12^. How can binding of Hsp70 to core-building stretches contribute to protein folding? (ii) Hsp70 acts with high on rate to compete with aggregation. How can slow-folding proteins complete their hydrophobic core in the presence of fast-rebinding Hsp70?

## Results

Looking for potentially analogous mechanisms in other chaperone systems, we note that the bacterial chaperonin GroEL-GroES first recruits substrates via a hydrophobic ring and subsequently exposes them to a larger, less hydrophobic chamber promoting folding of the enclosed intermediate (Fig. 1a). We hypothesize that Hsp70, offering a short, very hydrophobic cleft, and Hsp90, offering subsequently a large, less hydrophobic surface, may act as an analogous bipartite system providing an inverse hydrophobicity gradient along the folding path (Fig. 1b). Exposure to the Hsp90 surface may favor burial of hydrophobics within the core and thereby solve the Hsp70 paradoxes.

**Fig. 1.**
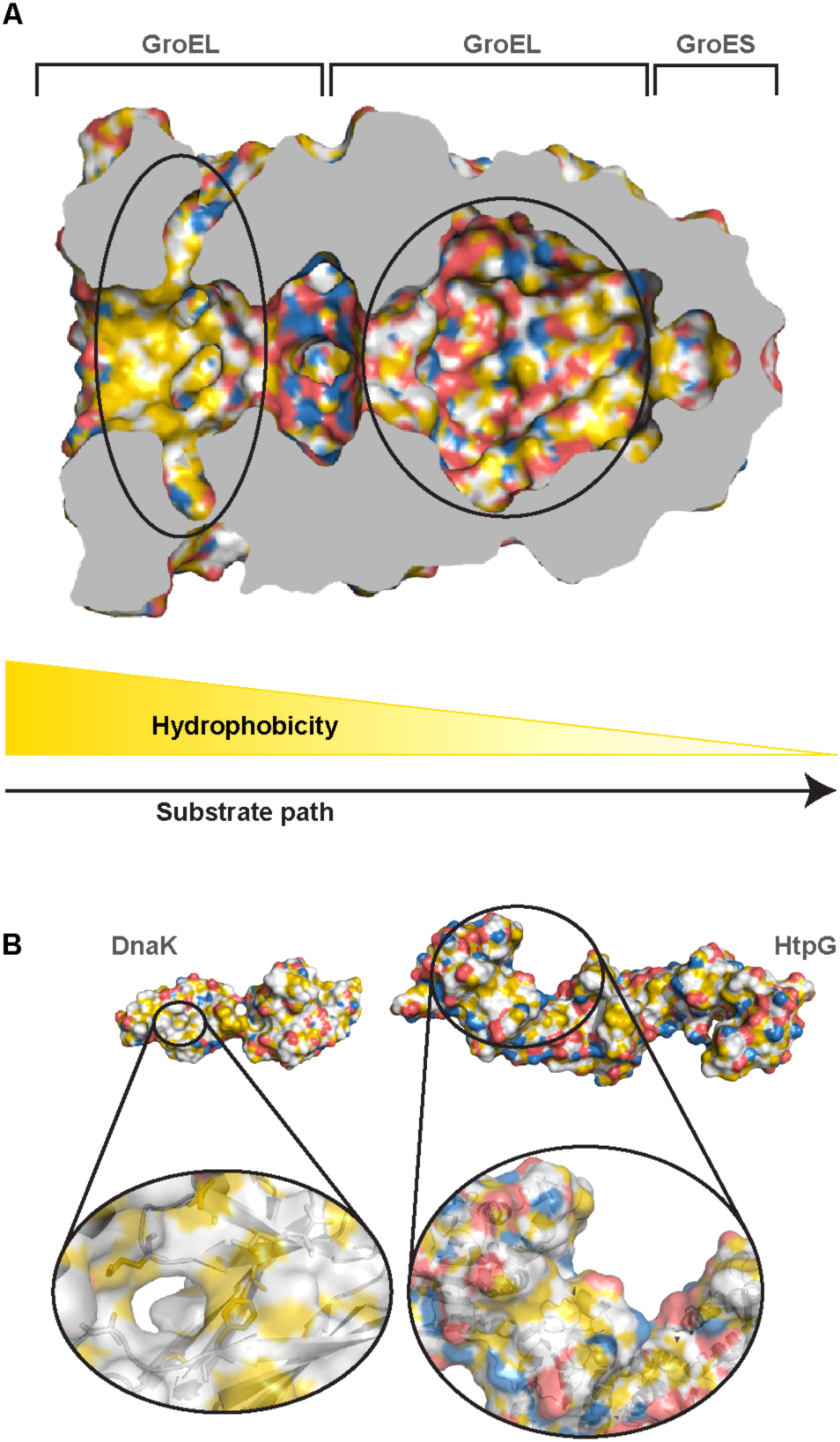
Hydrophobicity gradient in chaperone systems. YRB-coloured ^33^ chaperones highlight hydrophobicity and charge. **(A)** Cross-section of GroEL/GroES (pdb 1AON) shows the hydrophobic (yellow) entry chamber (left) and the more polar folding chamber (right). **(B)** The Hsp70-Hsp90 cascade offers a highly hydrophobic binding pocket on Hsp70 (pdb 2KHO) followed by a more polar surface on Hsp90.

Protein folding activity of Hsp70 chaperones was established both *in vivo* and *in vitro* using luciferase as a paradigmatic client, *nota bene* in the absence of Hsp90 ^11^. To address the Hsp70 paradoxes, we first revisited refolding of luciferase by the *E. coli* Hsp70 system, consisting of the Hsp70 DnaK, ATPase stimulating J-protein, DnaJ, and nucleotide exchange factor, GrpE. Substrate proteins binds to DnaK very fast, within seconds, but folding of e.g. luciferase takes around 30 mins ^11, 13, 14^. According to the second paradox, increasing Hsp70 levels should further favor the association of Hsp70 with unfolding protein, which may compete with productive folding ^15^. We, therefore, chemically denatured luciferase and monitored the refolding rate in the presence of the Hsp70 system (Fig. 2a). We kept concentration of luciferase, DnaJ and GrpE constant and titrated DnaK. We observed that independent of chaperone levels, activity of refolded luciferase reaches a plateau after around 30 min (Fig. 2b). The refolding yield, however, depended strongly on the concentration of Hsp70. It increased to a maximum of 82% refolded luciferase at 2 μM DnaK. Strikingly, however, increasing Hsp70 levels further successively reduced the yield, dropping to background levels at 10 μM DnaK (Fig. 2c). Hsp70 is thus not only a promoter but also an effective inhibitor of folding of luciferase.

**Fig. 2.**
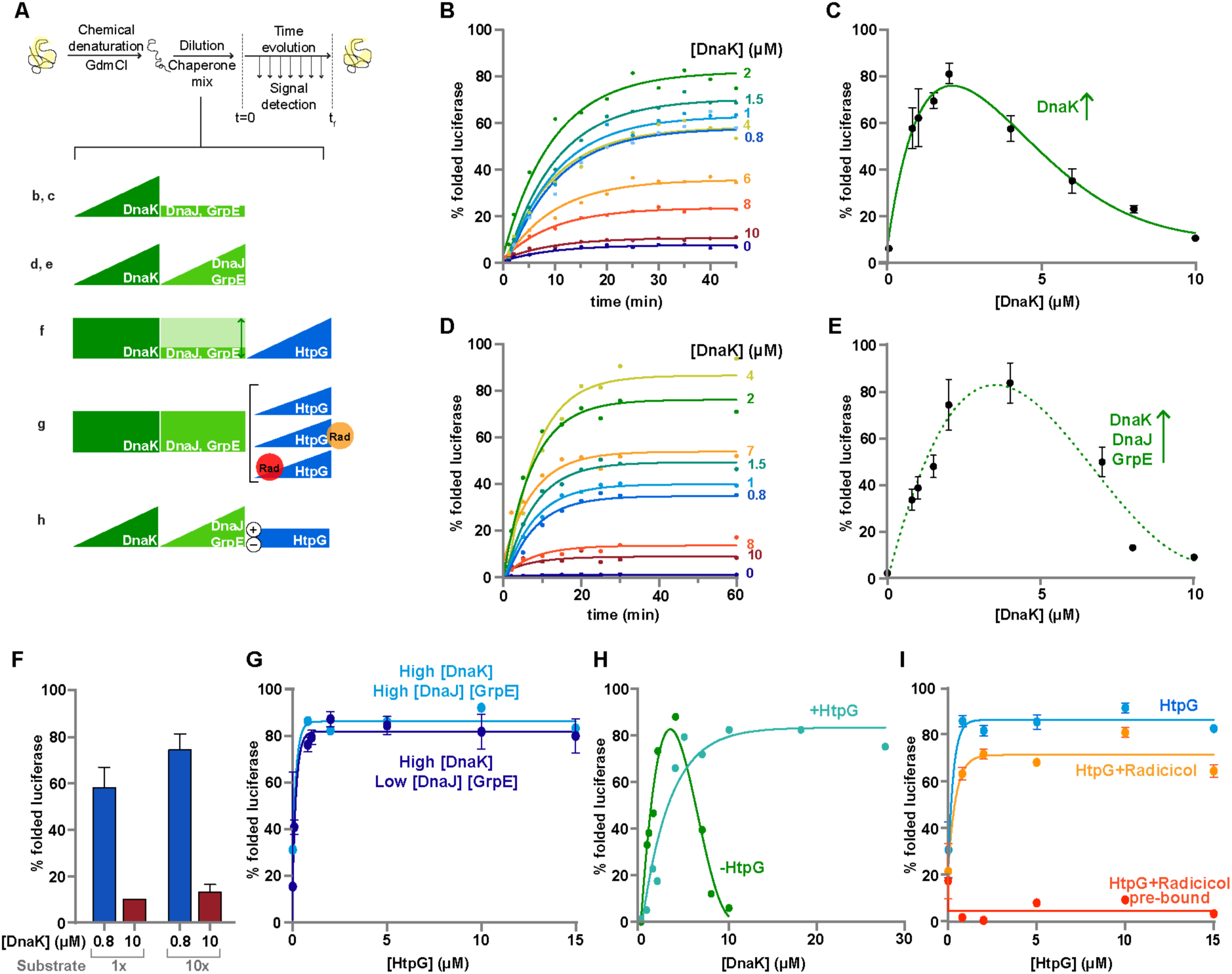
Hsp90 rescues luciferase out of the Hsp70 trap. **(A)** Experimental setup. **(B)** Time course of luciferase refolding in presence of constant levels of DnaJ (160 nM) and GrpE (400 nM) and increasing concentrations of DnaK (blue to red, 0 to 10 μM), fitted as first order reaction with identical rates. **(C)** High concentration of DnaK inhibits luciferase refolding. Plateau refolding yields in (B) plotted against DnaK concentration (±SEM). **(D)** Time course of luciferase refolding in presence of increasing DnaK/DnaJ/GrpE concentrations keeping a constant molar ratio (10:2:5). Fitting as in (B). **(E)** Plateau refolding yields in (D) plotted against DnaK concentration (±SEM). **(F)** Hsp70 folding block is independent of the chaperone/substrate ratio. Low DnaK concentration (blue, 0.8 μM) and high (red, 10 μM) show similar results at 80 nM or 800 nM of luciferase (±SEM). **(G)** Reactivation of denatured luciferase at high DnaK (blue) or DnaK/DnaJ/GrpE levels (cyan) as a function of HtpG concentration (±SD of plateau). **(H)** Titration of DnaK/DnaJ/GrpE keeping a constant ratio (10:2:5) in the absence (green) and presence (turquoise) of HtpG (1 μM). **(I)** ATPase activity of HtpG is required for substrate rescuing from DnaK. HtpG titration at high levels of DnaK/DnaJ/GrpE (10:2:5 μM; blue, no radicicol; orange, 30 μM radicicol; red, 30 μM radicicol pre-bound to Hsp90 (red) (±SD of plateau).

This phenomenon depends only on the levels of Hsp70 itself, and not on the relative ratio of chaperone to co-chaperones. Titrating DnaK, DnaJ and GrpE together revealed a similar picture, reaching a maximal refolding yield of 83.5% at 5 μM DnaK but subsequently falling to background levels at 10 μM (Fig. 2d, e). The physiological concentration of *E. coli* Hsp70 is even higher (∼ 27 μM at 30°C), and doubles upon heat shock ^16^. The Hsp70 block depends on the absolute Hsp70 levels, not on the ratio of Hsp70 to substrate. A 10-fold increase in luciferase levels did not release the Hsp70 block (Fig. 2f) Together, this suggests that Hsp70 requires an additional factor for effective refolding at physiological concentrations.

Given that Hsp90 acts downstream of Hsp70, we considered whether *E. coli* Hsp90, HtpG, restored folding activity at physiological Hsp70 levels ^7^. Hsp90 acts on steroid receptors downstream of Hsp70, and HtpG was found to have a mild effect on DnaK-dependent luciferase refolding ^9, 17–19^. Here we found that at high levels of the Hsp70 system (10 μM DnaK; Fig. 2c, e) Hsp90 restored its folding capacity, even when substoichiometric to DnaK (1 μM HtpG; Fig. 2g). Importantly, and in contrast to Hsp70, high levels of Hsp90 (15 μM) were not detrimental for folding. Thus, Hsp90 becomes essential for folding at high Hsp70 levels, without adverse side effects at physiological levels.

Next, we explored whether Hsp90 may function as safe guard to make the Hsp70 system robust to fluctuation in free chaperone levels, as naturally occurs upon and after cell stress. In the presence of Hsp90 (1 μM HtpG), we titrated the *E. coli* Hsp70 system up to physiological levels (27.4 μM DnaK / 1 μM DnaJ / 6.2 μM GrpE). The yield of refolded protein reaches a plateau above 5 μM DnaK (Fig. 2h). Thus, Hsp90 ensures that folding efficiency is independent of the levels of free Hsp70 and its co-chaperones.

To analyze the role of Hsp90 ATPase activity, we added the Hsp90-specific inhibitor Radicicol to the assay. Radicicol blocked Hsp90 activity in folding, consistent with earlier findings ^17^ (Fig. 2i). Remarkably, however, Radicicol only prevented substrate takeover by Hsp90 when pre-incubated with the chaperone. This implies that the Hsp90 ATPase is only relevant in the earliest phase on the folding path. As folding of luciferase continues for a further 30 min, we conclude that energy-dependent chaperone action takes place far before the transition state on the folding trajectory.

Next, we asked whether the interplay between the bacterial Hsp70 and Hsp90 systems is a conserved process. We confirmed that human Hsp70 is able to refold luciferase, assisted by the J-protein Hdj1 and the nucleotide exchange factor Apg2 (Fig. 3a) ^20^. We monitored luciferase refolding in the presence of increasing concentrations of all three members of the Hsp70 system (Fig. 3b, c). At 2 μM Hsp70 refolding of luciferase was maximal (75%). Further increase of Hsp70 subsequently reduced refolding efficiency, hitting basal refolding levels at 12 μM Hsp70. Addition of Hsp90, however, restored folding of luciferase at high concentrations in a substoichiometric manner (Fig. 3d). This indicates that the function of Hsp90 chaperones to counter the Hsp70-inflicted folding block is conserved between bacteria and man.

**Fig. 3.**
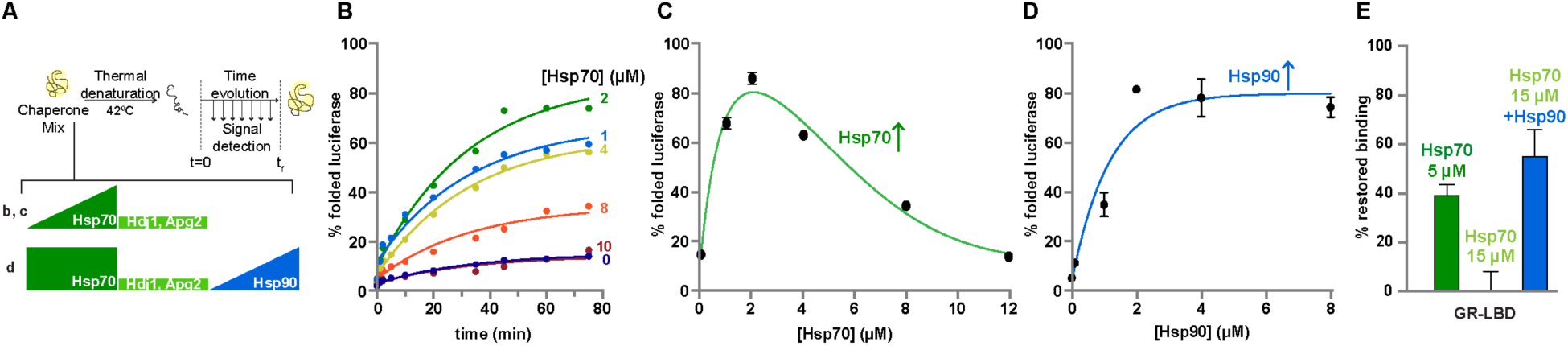
The effect of Hsp90 on the Hsp70 system is conserved and substrate independent. **(A)** Experimental setup of refolding experiments for thermally denatured luciferase or GR-LBD. **(B)** Time course of luciferase refolding at constant levels of Hdj1 (1 μM) and Apg2 (200 nM) and increasing Hsp70 (blue to red, 0 to 12 μM) fitted as first order reaction with identical rates. **(C)** High Hsp70 levels inhibit luciferase refolding. Plateau refolding yields plotted against Hsp70 concentration (±SD of plateau). **(D)** Hsp90 rescues luciferase refolding at high Hsp70 levels. Titration of Hsp90 and Hop keeping molar ratio (2:1) at constant high Hsp70/Hdj1/Apg2 levels (12:1:0.2 μM) (±SD of plateau). **(E)** Normalised recovered binding of F-Dex to GR-LBD after thermal unfolding with chaperone components indicated (± SD).

We now wondered how the assisted protein-folding path affected the folding rate of luciferase. Strikingly, neither presence nor concentration of any chaperone tested in this study does significantly influence the folding rate (Figs. 2bdgi and 3bd). In all experiments, the folding always followed a first order kinetics (averaged rates 5 ± 2 × 10^−4^ s^−1^ for 0-10 μM Hsp70 (Fig. 3b) and 6 ± 4 × 10^−4^ s^−1^ for 0-8 μM Hsp90 (Fig. 3d)). Therefore, the chaperones dramatically improve the refolding yield, but they do not change the refolding rate, and thus, not the transition state on the folding path. This is consistent with Anfinsen’s hypothesis for protein folding ^1^. We conclude that neither Hsp70 nor Hsp90 actively contribute to the folding process of luciferase. Thus, the Hsp70-Hsp90 cascade lacks foldase activity despite being required to generate high yields of refolded protein.

Next, we tested whether the mechanisms established here using luciferase could be confirmed for a classical Hsp90 *in vivo* and *in vitro* client, the ligand binding domain of the glucocorticoid receptor (GR-LBD) ^9, 19, 21^. Folding of GR-LBD can be monitored by binding to a fluorescent-labelled hormone derivative ^9, 21^. We thermally unfolded GR-LBD and monitored refolding at permissive temperature in the presence of Hsp70 and Hsp90. At high concentration of Hsp70 (15 μM), refolding stringently depended on Hsp90, in line with previous findings ^9, 19^. At low Hsp70 concentration (5 μM), however, Hsp90 was not essential for folding of GR-LBD (Fig. 3e). We conclude that the mechanistic interplay of Hsp70 and Hsp90 is also valid for folding of a protein that requires Hsp90 for maturation in the cell.

## Discussion

Together, our data demonstrate that Hsp90 has a conserved, general folding function downstream of Hsp70. Our findings provide a mechanistic explanation for earlier observations that presence of Hsp90 improves efficiency of the Hsp70 machine ^9, 10, 17, 18, 22, 23^. Combining current knowledge with the findings presented here, we propose a model for assisted protein folding in the cell according to a stop-start mechanism (Fig. 4a): I. A fraction of the protein folds spontaneously; II. The protein cycles on Hsp70, allowing the protein to re-enter pathway I, a step blocked by excess of Hsp70; III. Hsp90 takes over from Hsp70, promoting folding by breaking the deadlock of unproductive cycling. Therefore, there may be no functional need for repeated Hsp70 cycles of folding and release in the presence of Hsp90.

Our findings are consistent with the hydrophobic gradient hypothesis (Fig. 1b). Neither Hsp70 nor Hsp90 actively (re)fold the client (Fig. 4a). This is consistent with observations that the ATPase of DnaK is required for an early unfolding reaction, but not for subsequent folding ^24^. The Hsp70-Hsp90 cascade acts far ahead of the transition state of the folding reaction of the client protein. The task of the cascade is repositioning the folding client on the energy hypersurface such that it can ultimately fold autonomously according to Anfinsen, independent of chaperone action (Fig. 4b, c).

**Fig. 4.**
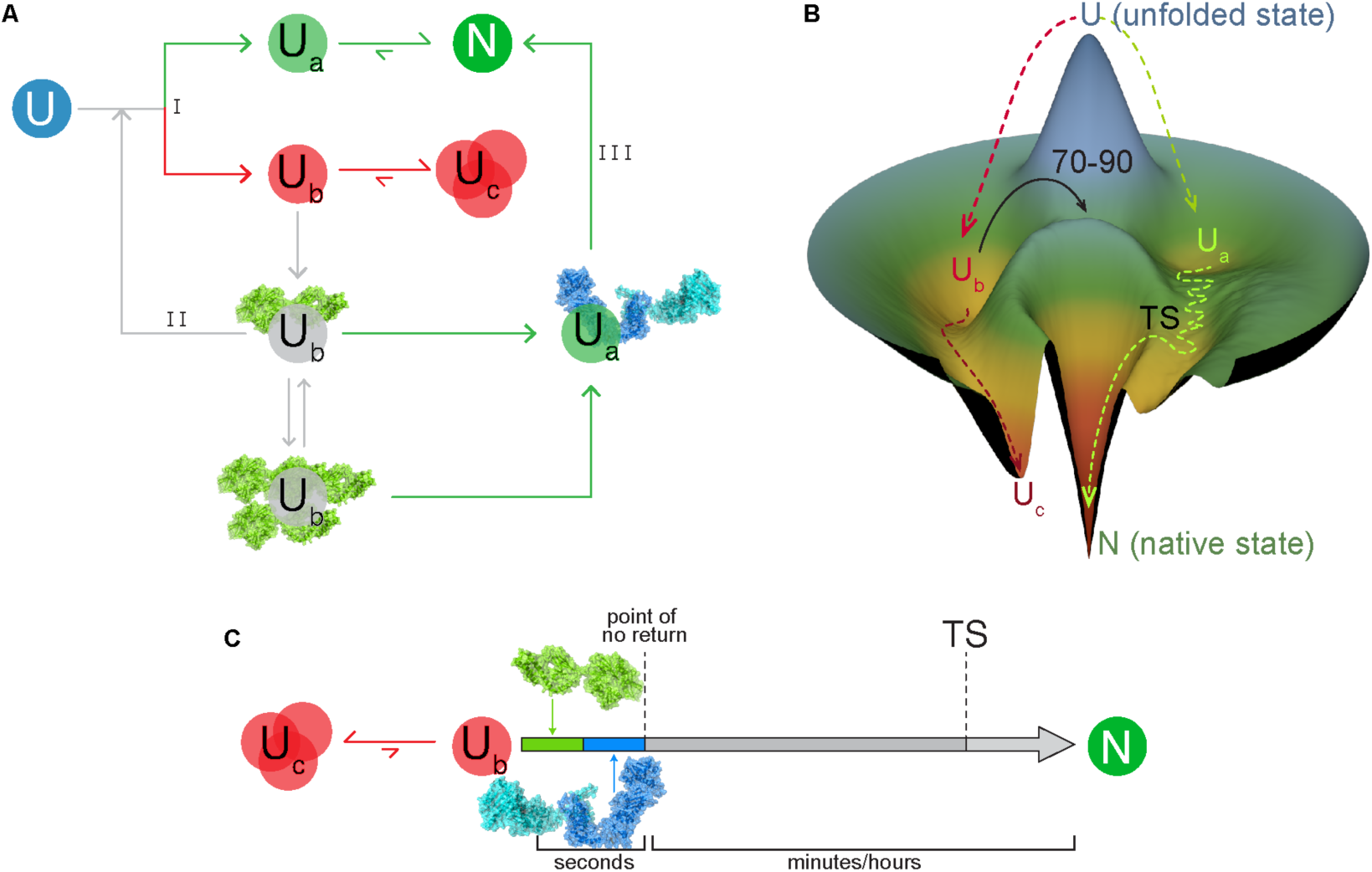
General model for chaperoned protein folding. **(A)** Chaperone assisted folding mechanism. Unfolded protein directly evolves to two states, folding competent U_a_ and aggregation/misfolding-prone U_b_, that binds to Hsp70. After release from Hsp70, the protein can further evolve to U_a_ or U_b_. However, high Hsp70 levels inhibit return to the U_a_/U_b_ junction. Hsp90 removes the Hsp70-inflicted folding block, promoting progression to the native state. Roman numbers indicate three modes of folding (I, spontaneous folding; II, Hsp70 cycling; III, Hsp70-Hsp90 cascade) **(B)** Three-dimensional scheme for protein folding. Energy landscape shows the effect of chaperones in early stages. U_b_ misfolded intermediate is boosted to a state that allows self-evolution following a slow path towards the native state. **(C)** Time-scheme indicating action of the Hsp70/Hsp90 cascade early on the folding path, far ahead of the transition state.

Components of the Hsp70-Hsp90 cascade are conserved in the cytosol from bacteria to man and are present in the main folding organelles of higher eukaryotes. This allows this cascade to take a major role in assisted protein folding throughout all kingdoms of life. The re-start activity of Hsp90 may be particularly crucial for slow folding, complex protein assemblies, making this chaperone essential in the cytoplasm of higher eukaryotes.

## Acknowledgments

We are grateful to Ineke Braakman for continuous support. We thank Friedrich Förster and Naomi Lodder for comments on the manuscript, Carlo van Mierlo, Daniel Fonseca and Doron Gollnast for discussion, Bastiaan Schravendeel for graphical 3D assistance and Elsbeth Schwarz for technical assistance. SGDR was supported by Marie-Curie Actions of the 7th Framework programme of the EU [Innovative Doctoral Programme “ManiFold” (No. 317371) and Initial Training Network “WntsApp” (No. 608180)], the Internationale Stichting Alzheimer Onderzoek (ISAO; project “Chaperoning Tau Aggregation”; No. 14542) and a ZonMW TOP grant (“Chaperoning Axonal Transport in neurodegenerative disease”; No. 91215084). This work was supported by the Deutsche Forschungsgemeinschaft (MA1278/4-3 to M.P.M.).

## Author contributions

SGDR and MPM conceived the study; SGDR, MPM and TML planned experiments; TML did experiments; SGDR, RK purified proteins and supported experiments; MPM and TML analyzed data; SGDR, MPM and TML wrote the manuscript.

## Materials and Methods

### Plasmids and protein purification

The bacterial protein **DnaK** was purified according to publish procedure ^14^. Briefly, DnaK was purified as native protein with an N-terminal His-SUMO fusion after overproduction in *ΔdnaK52* cells (BB1994). Upon expression, cell pellets were resuspendend in lysis buffer (20 mM Tris/HCl pH 7.9, 100 mM KCl, 1 mM PMSF) and subjected to lysis by microfluidizer EmulsiFlex C5 (Avestin, Ottawa, Canada) and afterwards applied onto a column with Ni-IDA resin (Macherey-Nagel, Düren, Germany). Subsequently, the column was washed with 20 CV of lysis buffer and 10 CV of ATP buffer (20 mM Tris/HCl pH 7.9, 100mM KCl, 5mM MgCl_2_, 5 mM ATP), and 2 CV of lysis buffer. The proteins were eluted with elution buffer (20 mM Tris/HCl pH 7.9, 100 mM KCl, 250 mM imidazol). To remove the His-SUMO tag from DnaK, the protein was treated with Ulp1 protease. Upon cleavage and dialysis into the lysis buffer, the protein mixture was subjected again to a Ni-IDA column and flow-through fraction containing tag-free DnaK was collected. Afterwards, DnaK was bound to anion exchange column (ResourceQ, GE Healthcare) equilibrated in low salt buffer (40 mM HEPES/KOH pH 7.6, 100 mM KCl, 5 MgCl_2_). DnaK was eluted with a linear KCl gradient (0.1–1 M) within 10 CV.

**DnaJ** was purified according to a published procedure ^25^ and is briefly described here. DnaJ was purified as native protein after overproduction in ΔdnaJ ΔcbpA cells. Upon expression, cell pellets were resuspended in lysis buffer (50 mM Tris/HCl pH 8, 10 mM DTT, 0.6% (w/v) Brij 58, 1 mM PMSF, 0.8 g/l Lysozyme) and lysed by microfluidizer EmulsiFlex-C5. Cell debris were removed by centrifugation at 20000 g for 30 min. 1 volume of buffer A (50 mM sodium phosphate buffer pH 7, 5 mM DTT, 1 mM EDTA, 0.1% (w/v) Brij 58) was added to the supernatant and DnaJ was precipitated by addition of (NH_4_)_2_SO_4_ to a final concentration of 65% (w/v). Upon centrifugation (15000 g, 30 min) the ammonium sulfate pellet was dissolved in 220 ml buffer B (50mM sodium phosphate buffer pH 7, 5 mM DTT, 1 mM EDTA, 0.1% (w/v) Brij 58, 2M Urea) and dialysed against the 5 l buffer B. Subsequently, DnaJ was loaded onto the cation exchange column (SP-sepharose, equilibrated with buffer B), washed with buffer B and eluted with a 15 CV long linear gradient of 0 to 666 mM KCl. DnaJ containing fractions were pooled and dialysed against 5 l buffer C (50 mM Tris/HCl, pH 7.5, 2 M urea, 0.1% (w/v) Brij 58, 5 mM DTT, 50 mM KCl). Afterwards the sample was loaded onto a hydroxyapatit column equilibrated in buffer C. The column was first washed with 1 CV buffer C supplemented with 1 M KCl, and then with 2 CV of buffer C. DnaJ was eluted with a linear gradient (0-50%, 1 CV) of buffer D (50 mM Tris/HCl, pH 7.5, 2 M urea, 0.1% (w/v) Brij 58, 5 mM DTT, 50 mM KCl, 600 mM KH_2_PO_4_) and 2 CV of 50% buffer D. The DnaJ containing fractions were pooled and dialysed against 2 l buffer E (50 mM Tris/HCl, pH 7.7, 100 mM KCl).

**GrpE** was purified as described before ^26^. Briefly, GrpE was purified after overproduction. Upon expression, cell pellets were resuspended in lysis buffer (Tris/HCl 50 mM, pH 7.5, 100 mM KCl, 3 mM EDTA, 1 mM PMSF) and lysed using a microfluidizer EmulsiFlex-C5. The lysate was clarified by centrifugation (18000 g, 50 min). Ammonium sulfate (0.35 g/ml) was added to the cleared lysate and insoluble proteins were separated from soluble by centrifugation (10000 g, 20 min). The pellet was dissolved in 200 ml buffer A (50 mM Tris/HCl pH 7.5, 100 mM KCl, 1 mM DTT, 1 mM EDTA, 10% glycerol) and dialysed twice against the same buffer (3 l, 4 h). Subsequently, protein was loaded onto anion exchange column (HiTrap Q XL; GE Healthcare) equilibrated with buffer A and GrpE was eluted using a linear gradient of buffer B (50 mM Tris/HCl pH 7.5, 1 M KCl, 1 mM DTT, 1 mM EDTA, 10% glycerol). Fractions containing the protein were dialyzed against buffer C (10 mM K_x_H_y_PO_4_ pH 6.8, 1 mM DTT, 10% glycerol). Next day the protein was loaded onto a Superdex 200 gel (GE Healthcare) filtration column equilibrated in buffer A, and concentrated using HiTrap Q XL with a steep gradient.

**HtpG** was purified as a native protein after overproduction in MC1061 cells induced by L-arabinose. Upon expression, cell pellets were resuspended in lysis buffer (25 mM K_x_H_y_PO_4_ pH 7.2, protease inhibitor (cOmplete™, EDTA free, Roche) and 5 mM ß-mercaptoethanol). The cells were lysed by microfluidizer EmulsiFlex-C5 and the lysate was clarified by centrifugation (20000 g, 40 min). The cleared lysate was loaded onto a Nickel Column (Poros 20MC) and eluted with elution buffer (25 mM K_x_H_y_PO_4_ pH 8, 400 mM NaCl, 500 mM imidazole, 5 mM ß-mercaptoethanol). Eluted fractions were diluted in lysis buffer and loaded onto an anion exchange colum (HiTrap Q XL). The protein was eluted from the column using high salt buffer (50 mM K_x_H_y_PO_4_ pH 7.2, 1 M KCl, 1mM ß-mercaptoethanol, 1 mM EDTA, 10% glycerol).

**Hdj1** was cloned and purified as a His_6_-SUMO fusion as described ^27^ using buffer (40 mM Hepes/KOH pH 7.6, 150 mM KCl, 5 mM MgCl_2_, 5% glycerol, 10 mM ß-mercaptoethanol) and Ni-IDA material.

**Hsp90α** was expressed and purified as described before ^28^. Briefly, Hsp90α was expressed from the bacterial expression vector pCA528 as fusion proteins with an N-terminal His_6_-Smt3 tag in the E. coli strain BL21(DE3) Star/pCodonPlus (Invitrogen). Upon expression, cells were resuspended in lysis buffer (40 mM Hepes/KOH pH 7.5, 100 mM KCl, 5 mM MgCl_2_, 10% glycerol, 4 mM β-mercaptoethanol, 5 mM PMSF, 1 mM Pepstatin A, 1 mM Leupeptin, and 1 mM Aprotinin) and lysed by a microfluidizer EmulsiFlex-C5. The lysate was clarified by centrifugation and incubated with Ni-IDA matrix. The matrix was loaded onto a column and washed with lysis buffer (without protease inhibitors) and eluted with lysis buffer containing 250 mM imidazole. Ulp1 was added to the eluted protein and the mixture was dialysed overnight against lysis buffer containing 20 mM KCl. Reverse Ni-IDA was applied to the dialysed protein followed by anion-exchange chromatography (ResourceQ™; GE healthcare) with a linear gradient of 0.02-1 M KCl. Fractions of eluted Hsp90 were subjected to Superdex 200 in storage buffer (40 mM Hepes/KOH pH 7.5, 50 mM KCl, 5 mM MgCl_2_, 10% glycerol, and 4 mM β-mercaptoethanol).

Human **Hsp70** was purified as native protein with an N-terminal His-SUMO fusion after overproduction in BL21 (DE3) Rosetta cells. After harvesting by centrifugation, pelleted cells were resuspended in lysis buffer (20 mM Tris/HCl pH 7.9, 100 mM KCl, 1 mM PMSF) and lysed using microfluidizer EmusiFlex C5. The lysate was clarified by centrifugation (20000 g, 50 min) and the supernatant was mixed with Ni-IDA beads and incubated for 15 min at 4°C. The lysate with beads was then poured into a column and was washed with 20 CV of lysis buffer (without PMSF), 20 CV of high salt buffer (20 mM Tris/HCl pH 7.9, 1 M KCl) and again with 2 CV of lysis buffer. The column was then slowly washed with 10 CV of ATP-buffer (40 mM Tris/HCl pH 7.9, 100 mM KCl, 5 mM MgCl_2_, 5 mM ATP) to elute bound substrates. Hsp70 was eluted with twice 1 CV of elution buffer (40 mM Tris/HCl pH 7.9, 100 mM KCl, 250 mM imidazole) and Ulp1 was added to the elution and dialyzed overnight against dialysis buffer (40 mM HEPES-KOH pH 7.6, 10 mM KCl, 5 mM MgCl_2_). The dialysed sample was then loaded on Ni-IDA material, flow-through containing Hsp70 was collected and loaded into a ResourceQ anion exchange column. Hsp70 was eluted with a linear gradient of AEX elution buffer (40 mM HEPES/KOH pH 7.6, 1 M KCl, 5 mM MgCl_2_, 10 mM ß-mercaptoethanol, 5% glycerol) and dialysed against storage buffer (40 mM HEPES/KOH pH 7.6, 50 mM KCl, 5 mM MgCl_2_, 10 mM ß-mercaptoethanol, 10% glycerol).

**Hop** and **Apg2** were purified as native proteins both with an N-terminal His-SUMO fusion after overproduction in BL21 (DE3) Rosetta cells (Merck KGaA, Darmstadt, Germany). Upon expression, the cell pellets were resuspended in lysis buffer (40 mM Tris/HCl pH 7.9, 100 mM KCl, 5 mM ATP, 8 mg/l Pepstatin, 10 μg/ml Aprotinin, 5 mg/l Leupeptin). The cells were lysed by microfluidizer EmulsiFlex C5. The lysate was clarified by centrifugation (20000 g, 40 min). Ni-IDA resin was added to the cleared lysate and incubated for 20 min, and then washed with 20 CV wash buffer (40 mM Tris/HCl pH 7.9, 100 mM KCl, 5 mM ATP) followed by elution with elution buffer (40 mM Tris/HCl pH 7.9, 100 mM KCl, 300 mM imidazol). Subsequently, buffer exchange was performed on HiPrep 26/10 Desalting column (GE Healthcare) equilibrated in the desalting buffer (20 mM Tris/HCl pH 7.9, 100 mM KCl, 10% glycerol). Upon buffer exchange, to remove the His-SUMO-tag, the protein was incubated overnight at 4 °C with Ulp1 protease in the presence of 5 mM ATP. Next day the protein was loaded onto HiLoad 16/600 Superdex 200 column equilibrated in gel filtration buffer (40 mM HEPES/KOH pH 7.6, 10 mM KCl, 5 mM MgCl2, 10% glycerol). Protein containing fractions were pooled and subjected to anion-exchange chromatography on ResourceQ column (GE Healthcare) for Apg2, and POROS 20HQ column (Thermo Fisher Scientific) in the case of Hop, both equilibrated with gel filtration buffer. In both cases the protein was eluted with linear gradient KCl gradient (0.01–1 M) within 10 CV.

**Firefly luciferase** was purified according to previous procedures ^20^. Briefly, firefly luciferase was expressed in XL10 Gold^®^ strain (Stratagene, US). Cells containing the expression plasmid were grown at 37°C until OD_600_ = 0.5 was reached, at which point the temperature was lowered to 20°C. After 45 minutes shaking at 20°C, cells were induced with IPTG overnight. Upon harvesting by centrifugation, pellets were resuspended in precooled lysis buffer (50 mM Na_x_H_y_PO_4_ pH 8.0, 300 mM NaCl, 10 mM ß-mercaptoethanol, protease inhibitor (cOmplete™, EDTA free, Roche) DNase) and lysed using a microfluidizer EmulsiFlex-C5. The lysate was cleared and incubated with Ni-IDA resin for 30 min. Subsequently, the lysate-protino mixture was loaded onto a column and washed with 10 CV of lysis buffer, 10 CV of wash buffer (50 mM Na_x_H_y_PO_4_ pH 8.0, 300 mM NaCl and 10 mM ß-mercaptoethanol) and eluted by addition of elution buffer (50 mM Na_x_H_y_PO_4_ pH 8.0, 300 mM NaCl, 250 mM Imidazole, 5 mM ß-mercaptoethanol) collecting 1-2 ml fractions. Luciferase was dialyzed overnight using dialysis buffer (50 mM Na_x_H_y_PO_4_ pH 8.0, 300 mM NaCl and 10 mM ß-mercaptoethanol, 10% glycerol).

**Ydj1** was overproduced by addition of IPTG and cells were grown for additional 5 h. Cells were harvested by centrifugation, washed twice with H_2_O and resuspended in a minimal volume of 1x lysis buffer (10x lysis buffer: 180 mM Spermidin/HCl pH 7.6, 1 M (NH_4_)_2_SO_4_, 50 mM DTT, 5 mM EDTA) in buffer A (50 mM Tris/HCl, pH 7.6, 10% (w/w) saccharose, 1 mM PMSF). Cells were disrupted in a microfluidizer EmulsiFlex-C5 and cell debris removed by centrifugation. (NH_4_)_2_SO_4_ was added to the clarified lysate to a saturation of 85% over a 30 min period, and stirring continued for additional 30 min at 4°C. Ydj1 was separated from precipitated proteins by centrifugation (20 min, 15000 g) and dialyzed overnight against Ydj1 buffer (HEPES/KOH 40 mM pH 7.6, 150 mM KCl, 1 mM DTT, 5% glycerol). The protein was then loaded onto an anion exchange column (DEAE-Sepharose), washed with Ydj1 buffer and eluted with a linear gradient of 0-800 mM KCl over 3 CV. The fractions containing Ydj1 were collected, concentrated and loaded onto a Superdex 200™ and subsequently loaded onto a ResourceQ™ column for further purification and concentration. Ydj1 was stored in Ydj1 buffer.

For **GR-LBD**, the protocol previously described ^28^ was followed. Briefly, the GR-LBD fragment (521-777) with an F602S amino acid exchange was expressed in BL21 Star (DE3) Rosetta. Cells containing the expression plasmid were grown at 37°C until OD_600_ = 0.8 was reached. Dexamethasone was then added to a final concentration of 250 μM and cells were induced with 0.5 mM IPTG at 18°C overnight. Cells were lysed using a microfluidizer EmulsiFlex-C5 in MBP-lysis buffer (50 mM Tris HCl pH 8.3, 300 mM KCl, 5 mM MgCl_2_, 0.04% CHAPS, 1 mM EDTA, 10% glycerol, 50 μM dexamethasone, 5 mM ß-mercaptoethanol) with protease inhibitor tablets (Roche) and purified using an amylose resin (NEB, E8021S) following the manufacturer protocol with 50 mM dexamethasone addition to every buffer. To remove dexamethasone the eluted protein was dialysed in 2 l MBP-lysis buffer (without dexamethasone) for four times, each at least 2 h. Protein was freshly purified before use.

### Luciferase refolding assays

For the bacterial system, firefly luciferase (10 μM) was chemically denatured by incubation in unfolding buffer (5 M GdmCl, 30 mM Tris/acetate pH 7.5) for 10 min at 22°C. For refolding, luciferase was diluted 125-fold into refolding buffer (25 mM HEPES/KOH pH 7.6, 100 mM KOAc, 10 mM Mg(OAc)_2_, 2 mM ATP, 5 mM DTT; 80 nM final luciferase concentration) containing the specified chaperone concentrations and incubated at 30°C. Luciferase activity was determined in a Lumat LB 9507 luminometer (Berthold Technologies) by transferring 1 μl of sample to 124 μl of assay buffer (100 mM phosphate buffer pH 7.6, 25 mM Glycylglycine, 100 mM KOAc, 15 mM Mg(OAc)_2_, 5 mM ATP) at the indicated time points. 125 μl of 0.1 μM luciferin were injected by the instrument right before each measurement and luminescence was measured for 5 s.

For the human system, 80 nM firefly luciferase in refolding buffer containing the indicated amounts of chaperones was heat-denatured at 42°C for 10 min. At the indicated time points, 1 μL sample was diluted into 124 μl of assay buffer and measured as described for the bacterial system. Refolding yields were normalized based on the activity of non-denatured luciferase.

### Luciferase data fitting

Data fitting was done using Prism 6 (GraphPad software, USA).

In case of **Hsp70 titrations**, <u>% refolding *vs* time</u> data was fitted to a one-phase association equation (Figure 1b, 1d, 2b) with a shared *k value,* corresponding to a first order reaction:

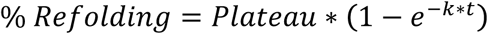

**Table 1.** K values and their standard error from the fit of data in figure 1b, 1d and 2b to a one phase association equation.

|  | K value | ±SE |
| --- | --- | --- |
| Fig. 1b | 0,093 | 0,004 |
| Fig. 1d | 0,115 | 0,006 |
| Fig. 2b | 0,033 | 0,002 |

For each concentration, the percentage of refolded luciferase at the plateau were extrapolated from the fits and standard error of the fit for the plateau were calculated:

**Table 2.** Plateau values and their standard error from the fit of data in figure 1b, 1d and 2b.

| Fig. 1b |  |  |
| --- | --- | --- |
| Conc | Plateau | ±SE |
| 0 | 7,35 | 0,27 |
| 0.8 | 59,66 | 1,51 |
| 1 | 64,26 | 1,42 |
| 1.5 | 71,77 | 1,11 |
| 2 | 83,49 | 1,79 |
| 4 | 59,28 | 1,21 |
| 6 | 36,14 | 0,81 |
| 8 | 23,67 | 0,42 |
| 10 | 10,79 | 0,26 |

| Fig. 1d |  |  |
| --- | --- | --- |
| Conc | Plateau | ±SE |
| 0 | 1,01 | 0,03 |
| 0.8 | 35,73 | 1,38 |
| 1 | 40,88 | 0,99 |
| 1.5 | 50,49 | 1,09 |
| 2 | 77,80 | 1,53 |
| 4 | 89,14 | 3,75 |
| 7 | 52,53 | 1,58 |
| 8 | 13,78 | 0,77 |
| 10 | 8,18 | 0,71 |

| Fig. 2b |  |  |
| --- | --- | --- |
| Conc | Plateau | ±SE |
| 0 | 14,36 | 0,29 |
| 1 | 66,04 | 2,34 |
| 2 | 83,41 | 2,36 |
| 4 | 61,09 | 1,08 |
| 8 | 33,36 | 1,45 |
| 12 | 13,50 | 1,09 |

<u>% Refolding *vs* [Hsp70]</u> data (Fig. 1c and 2c) fitted accurately to an equation derived from the chemical mechanisms of the reaction implying that more than one Hsp70 molecule binds to one substrate and binding of two Hsp70 irreversibly changes the substrate:

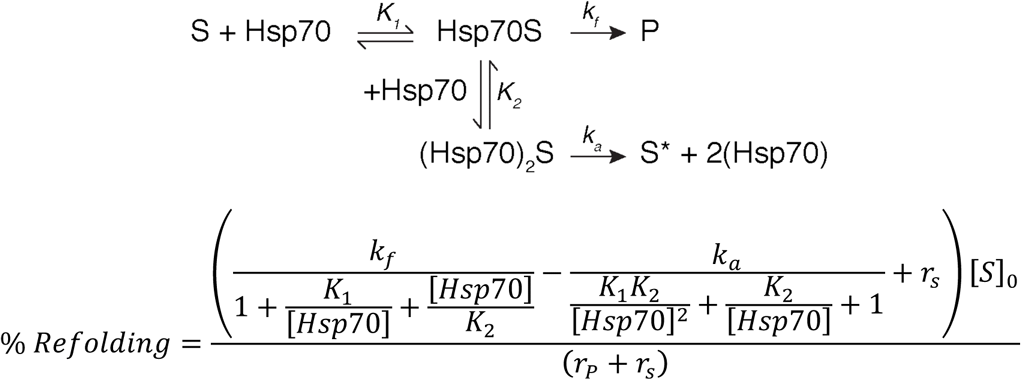

**Equation 1.** Equation derived from the mechanistic scheme of the chemical reaction shown above. r_p_ represents the observed rate of the individual refolding reactions in Fig. 1b and 1d, whilst r_s_ is the rate of the competing reaction. The values for all the parameters of the fit can be found in the following table, including the constrained *S* and *r*_*p*._

**Table 3.**
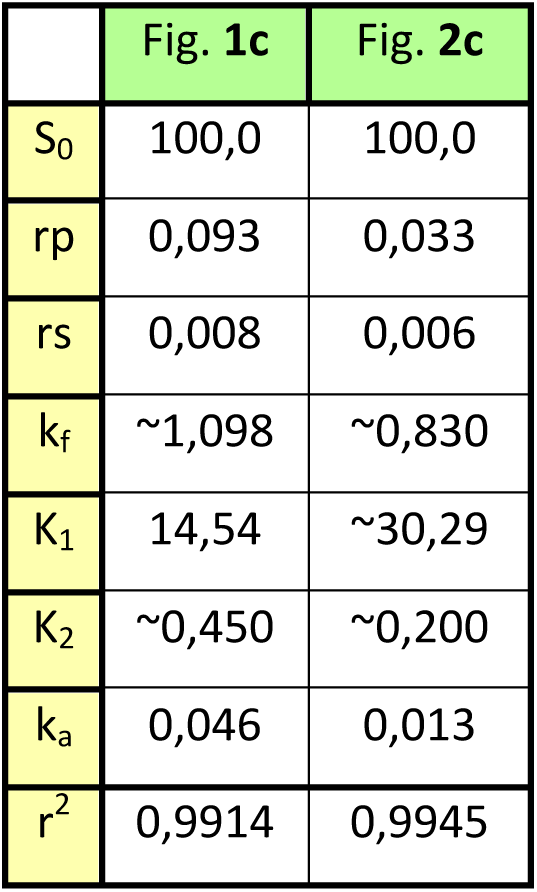
Values of the parameters of the derived equation used in Fig. 1c and 1d.

|  | Fig. 1c | Fig. 2c |
| --- | --- | --- |
| $S_0$ | 100,0 | 100,0 |
| $r_p$ | 0,093 | 0,033 |
| $r_s$ | 0,008 | 0,006 |
| $k_f$ | ~1,098 | ~0,830 |
| $K_1$ | 14,54 | ~30,29 |
| $K_2$ | ~0,450 | ~0,200 |
| $k_a$ | 0,046 | 0,013 |
| $r^2$ | 0,9914 | 0,9945 |

*Equation 1* assumes Hsp70 concentration as constant since it is in large excess over the substrate. In the situation of Fig. 1e, not only Hsp70 concentration changes but its co-chaperones are titrated up as well. DnaJ was shown to bind as a substrate to DnaK when no substrate is present or, presumably, when in large excess over the substrate ^29^. That translates in a more complex situation that could not be described by the same equation. In this case, a dotted line guides the eye.

For the **Hsp90 titrations**, <u>% refolding *vs* time</u> data was fitted to a one-phase association rate equation as described for the Hsp70 experiments with a shared k value (graphs not shown). <u>Refolding *vs* [Hsp90]</u> was fitted to a one-phase association equation with association constant K (Figure 1g, 1i, 2d).

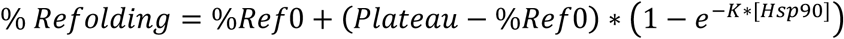

**Table 4.**
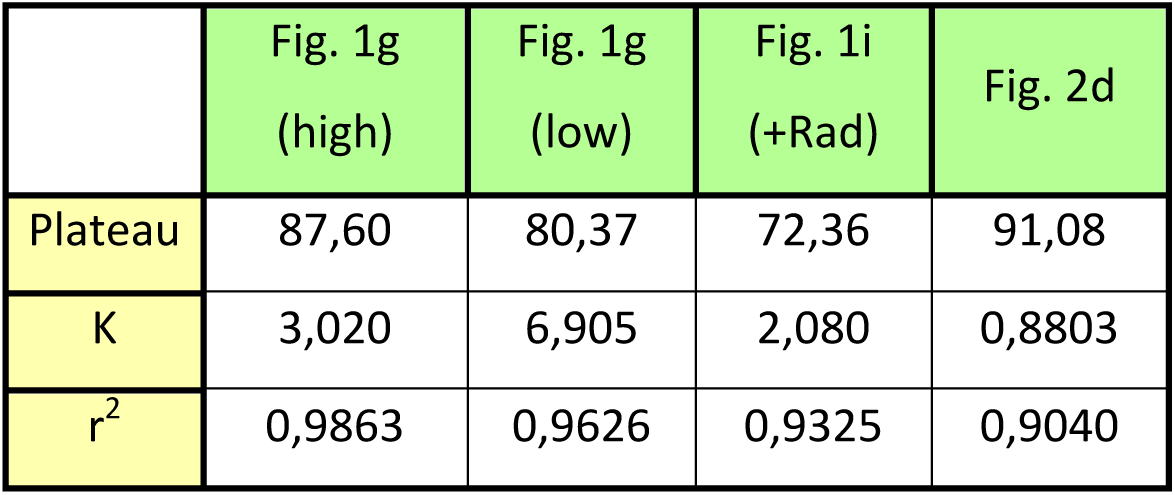
Values of the parameters of the one phase association equation used to fit data in Fig. 1g, 1i and 2d (Hsp90 titration experiments).

### GR-LBD binding assay

2 μM GR-LBD was denatured in the absence or presence of 15 μM Hsp70, 2 μM Ydj1, 2 mM ATP and 5 mM MgCl_2_ at 42°C for 10 minutes. After denaturation when indicated, 15 μM Apg2 and 15 μM Hsp90 were added. Fluorescence anisotropy was measured on a plate reader (CLARIOStar, BMG Labtech) with excitation/emission wavelengths of 485/538 nm. Ligand binding was initiated by addition of F-Dex (fluorescein-labelled dexamethasone) to a final concentration of 20 nM and association was monitored measuring fluorescence anisotropy over time until reaching saturation. Control samples with GR-LBD alone ± Hsp70 were prepared with matching volumes of the chaperones storage buffer. Normalization was done considering 100% and 0% binding of F-Dex as folded and unfolded GR-LBD respectively.

### PyMOL Structures

GroEL/GroES (pdb file: 1AON) ^30^, DnaK (pdb file: 2KHO) ^31^, HtpG ^32^ are colored using YRB highlighting scheme in PyMOL ^33^.

## References

[1] Anfinsen, C. B. Principles that govern the folding of protein chains. Science 181, 223–230 (1973).

[2] Daggett, V. & Fersht, A. The present view of the mechanism of protein folding. Nat. Rev. Mol. Cell Biol. 4, 497–502 (2003).

[3] Kim, Y. E., Hipp, M. S., Bracher, A., Hayer-Hartl, M. & Hartl, F. U. Molecular chaperone functions in protein folding and proteostasis. Annu. Rev. Biochem. 82, 323–355 (2013).

[4] Buchberger, A., Bukau, B. & Sommer, T. Protein quality control in the cytosol and the endoplasmic reticulum: brothers in arms. Mol. Cell 40, 238–252 (2010).

[5] Ellis, J. Proteins as molecular chaperones. Nature 328, 378–379 (1987).

[6] Mayer, M. P. Hsp70 chaperone dynamics and molecular mechanism. Trends Biochem. Sci. 38, 507–514 (2013).

[7] Karagöz, G. E. & Rüdiger, S. G. D. Hsp90 interaction with clients. Trends Biochem. Sci. 40, 117–125 (2015).

[8] Karagöz, G. E. et al. Hsp90-Tau complex reveals molecular basis for specificity in chaperone action. Cell 156, 963–974 (2014).

[9] Kirschke, E., Goswami, D., Southworth, D., Griffin, P. R. & Agard, D. A. Glucocorticoid receptor function regulated by coordinated action of the Hsp90 and Hsp70 chaperone cycles. Cell 157, 1685–1697 (2014).

[10] Wegele, H., Wandinger, S. K., Schmid, A. B., Reinstein, J. & Buchner, J. Substrate transfer from the chaperone Hsp70 to Hsp90. J. Mol. Biol. 356, 802–811 (2006).

[11] Schröder, H., Langer, T., Hartl, F. U. & Bukau, B. DnaK, DnaJ and GrpE form a cellular chaperone machinery capable of repairing heat-induced protein damage. EMBO J. 12, 4137–4144 (1993).

[12] Rüdiger, S., Germeroth, L., Schneider-Mergener, J. & Bukau, B. Substrate specificity of the DnaK chaperone determined by screening cellulose-bound peptide libraries. Embo J 16, 1501–7 (1997).

[13] Hu, B., Mayer, M. P. & Tomita, M. Modeling Hsp70-mediated protein folding. Biophys. J. 91, 496–507 (2006).

[14] Kityk, R., Vogel, M., Schlecht, R., Bukau, B. & Mayer, M. P. Pathways of allosteric regulation in Hsp70 chaperones. Nat. Commun. 6, 8308 (2015).

[15] Kellner, R. et al. Single-molecule spectroscopy reveals chaperone-mediated expansion of substrate protein. Proc. Natl. Acad. Sci. U. S. A. 111, 13355–13360 (2014).

[16] Mogk, A. et al. Identification of thermolabile Escherichia coli proteins: prevention and reversion of aggregation by DnaK and ClpB. Embo J 18, 6934–49 (1999).

[17] Genest, O., Hoskins, J. R., Camberg, J. L., Doyle, S. M. & Wickner, S. Heat shock protein 90 from Escherichia coli collaborates with the DnaK chaperone system in client protein remodeling. Proc. Natl. Acad. Sci. U. S. A. 108, 8206–8211 (2011).

[18] Genest, O., Hoskins, J. R., Kravats, A. N., Doyle, S. M. & Wickner, S. Hsp70 and Hsp90 of E. coli Directly Interact for Collaboration in Protein Remodeling. J. Mol. Biol. 427, 3877–3889 (2015).

[19] Sanchez, E. R. et al. Relationship of the 90-kDa murine heat shock protein to the untransformed and transformed states of the L cell glucocorticoid receptor. The Journal of biological chemistry 262, 6986–6991 (1987).

[20] Rampelt, H. et al. Metazoan Hsp70 machines use Hsp110 to power protein disaggregation. EMBO J. 31, 4221–4235 (2012).

[21] Lorenz, O. R. et al. Modulation of the Hsp90 chaperone cycle by a stringent client protein. Mol. Cell 53, 941–953 (2014).

[22] Schumacher, R. J. et al. Cooperative action of Hsp70, Hsp90, and DnaJ proteins in protein renaturation. Biochemistry (N. Y.) 35, 14889–14898 (1996).

[23] Nakamoto, H. et al. Physical interaction between bacterial heat shock protein (Hsp) 90 and Hsp70 chaperones mediates their cooperative action to refold denatured proteins. J. Biol. Chem. 289, 6110–6119 (2014).

[24] Sharma, S. K., De los Rios, P., Christen, P., Lustig, A. & Goloubinoff, P. The kinetic parameters and energy cost of the Hsp70 chaperone as a polypeptide unfoldase. Nat. Chem. Biol. 6, 914–920 (2010).

[25] Graf, C., Stankiewicz, M., Kramer, G. & Mayer, M. P. Spatially and kinetically resolved changes in the conformational dynamics of the Hsp90 chaperone machine. EMBO J. 28, 602–613 (2009).

[26] Schönfeld, H. J., Schmidt, D., Schröder, H. & Bukau, B. The DnaK chaperone system of Escherichia coli: quaternary structures and interactions of the DnaK and GrpE components. J. Biol. Chem. 270, 2183–2189 (1995).

[27] Malakhov, M. P. et al. SUMO fusions and SUMO-specific protease for efficient expression and purification of proteins. J. Struct. Funct. Genomics 5, 75–86 (2004).

[28] Nguyen, M. T. et al. Isoform-Specific Phosphorylation in Human Hsp90beta Affects Interaction with Clients and the Cochaperone Cdc37. J. Mol. Biol. 429, 732–752 (2017).

[29] Laufen, T. et al. Mechanism of regulation of hsp70 chaperones by DnaJ cochaperones. Proc. Natl. Acad. Sci. U. S. A. 96, 5452–5457 (1999).

[30] Xu, Z., Horwich, A. L. & Sigler, P. B. The crystal structure of the asymmetric GroEL-GroES-(ADP)7 chaperonin complex. Nature 388, 741–750 (1997).

[31] Bertelsen, E. B., Chang, L., Gestwicki, J. E. & Zuiderweg, E. R. Solution conformation of wild-type E. coli Hsp70 (DnaK) chaperone complexed with ADP and substrate. Proc. Natl. Acad. Sci. U. S. A. 106, 8471–8476 (2009).

[32] Krukenberg, K. A., Street, T. O., Lavery, L. A. & Agard, D. A. Conformational dynamics of the molecular chaperone Hsp90. Q. Rev. Biophys. 44, 229–255 (2011).

[33] Hagemans, D., van Belzen, I. A., Morán Luengo, T. & Rüdiger, S. G. D. A script to highlight hydrophobicity and charge on protein surfaces. Front. Mol. Biosci. 2, 56 (2015).

